# Mixed outcomes in recombination rates after domestication: Revisiting theory and data

**DOI:** 10.1101/2024.08.07.607048

**Authors:** Madeline Bursell, Manav Rohilla, Lucia Ramirez, Yuhuan Cheng, Enrique J. Schwarzkopf, Rafael F. Guerrero, Caiti Smukowski Heil

## Abstract

The process of domestication has altered many phenotypes. Selection on these phenotypes has long been hypothesized to indirectly select for increases in recombination rate. This hypothesis is consistent with theory on the evolution of recombination rate, but empirical support has been unclear. We review relevant theory, lab-based experiments, and data comparing recombination rates in wild progenitors and their domesticated counterparts. We utilize population sequencing data and a deep learning method to infer genome-wide recombination rates for new comparisons of chicken/red junglefowl, sheep/mouflon, and goat/bezoar. We find evidence of increased recombination in domestic goats compared to bezoars, but more mixed results in chicken, and generally decreased recombination in domesticated sheep compared to mouflon. Our results add to a growing body of literature in plants and animals that finds no consistent evidence of an increase in recombination with domestication.

## Introduction

Intentional and unintentional selection on traits has shaped numerous features of modern day crops, animals, and microorganisms. Generally, this process of domestication is envisioned to begin with an initial stage of cultivation of a wild species followed by a recent phase of more intense artificial selection for breed/line formation. In addition to changes in phenotypes of selected traits, domestication has resulted in changes including increased ploidy or a switch of mating systems in some species, and common hallmarks like a reduction in effective population size, an increase in deleterious variants, reduced genetic diversity, and increased homozygosity (Moyers et al., 2018).

Domestication is also hypothesized to alter meiotic recombination landscapes in domesticated species compared to their wild counterparts. Meiotic recombination describes the reciprocal exchange of genetic material between homologous chromosomes, and is required for the successful completion of meiosis in most sexually reproducing organisms. Errors in recombination are associated with aneuploidy and loss of fertility (Ariad et al., 2024; Fledel-Alon et al., 2009; Hassold & Hunt, 2001; Roeder, 1997). Apart from its mechanistic role in meiosis, recombination is important in shuffling alleles between homologous chromosomes, thus increasing genetic diversity and increasing the efficacy of selection in populations. Recombination rate varies between species, populations, sexes, individuals, and across individual genomes (Smukowski & Noor, 2011; Stapley Jessica et al., 2017). A minimum number of one crossover per chromosome (or chromosome arm) is postulated to set a minimum threshold for recombination rate (F & C, 2001; Lynch, 2006). The dramatic variation that exists above this threshold has fascinated many researchers in the field of evolutionary biology for decades.

Numerous hypotheses have been put forward to explain the evolution of recombination rate, including meiotic drive, karyotype changes, and selection directly on recombination rates, or indirectly due to adaptation to new or fluctuating environments, host-parasite interactions, or directional selection on a phenotype (Arter & Keeney, 2023; Bomblies et al., 2015; Dapper & Payseur, 2017; Henderson & Bomblies, 2021; Otto, 2009; Otto & Payseur, 2019; Ritz et al., 2017). The hypothesis that domestication can increase recombination rates is based on the latter, that repeated directional selection on a trait, or suite of traits, can indirectly select for higher recombination rates to increase the efficacy of selection. We start by reviewing the theoretical contributions to this hypothesis, then review lab based directional selection experiments, comparisons between recombination landscapes of domesticated and wild populations, and add new comparisons of sheep/mouflon, goat/bezoar, and chicken/junglefowl recombination landscapes. Overall, we find evidence of increased, decreased, and unchanged recombination rates between wild and domesticated organisms.

### Theoretical basis of the domestication theory

The evolution of sex and recombination has a rich theoretical history. Here, we will focus our discussion on theory motivating the domestication hypothesis, but for more thorough review and discussion see (Dapper & Payseur, 2017; de Visser & Elena, 2007; Feldman et al., 1996; Otto, 2009; Otto & Gerstein, 2006; Rice, 2002). Models suggest that recombination rate can increase when a recombination modifier is linked to two or more alleles under selection that are on different haplotypes more often than expected by chance (negative linkage disequilibrium). In infinite populations, there is a narrow parameter space when epistasis is weak and negative, under which recombination modifiers can increase recombination (Barton, 1995; Charlesworth, 1993). Because of the requirements of universally negative epistasis, these models have not been thought to fit empirical data well. Instead, the field has coalesced around models where increased recombination rate can evolve to reduce selective interference between loci, dubbed Hill-Robertson interference (Barton & Otto, 2005; Felsenstein & Yokoyama, 1976; Hill & Robertson, 1966; Otto & Barton, 2001).

Briefly, when new beneficial alleles arise in a finite population, they may occur on a background with other beneficial alleles (positive disequilibrium) or on a background with deleterious alleles (negative disequilibrium). With negative disequilibrium, genetic variation is hidden from selection by beneficial mutations being linked to deleterious mutations, slowing selection as these chromosome combinations compete with each other (i.e. interference). But if recombination is increased by a recombination modifier that is linked to loci under selection, beneficial alleles can be combined onto the same haplotype and the linked modifier will spread with the favored haplotype. Selection on recombination rate modifiers will be greatest when multiple linked loci are repeatedly under selection, and increased recombination can evolve with moderate to strong selection on loci linked to the modifier (Otto & Barton, 2001). Moreover, models based on this foundation of selection in finite populations, which we’ll refer to as drift-based models, are fairly robust to the presence and sign of epistasis and the type of selection, and extend to large populations if enough loci are under selection or if the population is spatially structured (Iles et al., 2003; Keightley & Otto, 2006; Martin et al., 2006).

This scenario is compatible with our framework of domestication, and domestication has been considered a plausible empirical test for the drift-based hypothesis of the evolution of recombination. This theory predicts that domesticated organisms should have increased recombination rates compared to their wild progenitors. While these models dictate that the recombination modifier must be linked to the loci under selection and are concerned only with local recombination (i.e. only between the loci under selection), it is conceivable that modifiers that evolve through their local effects lead to an overall increase in recombination across the genome.

### Empirical studies of directional selection and recombination rate in the lab

A number of experiments have been carried out in the lab that test various features and predictions of the drift-based model for the evolution of recombination rate, offering a simplified view of the forces at play in the more complicated case of domestication. The base assumption for recombination rate to evolve is that recombination rate must show heritable variation in a population. It is now well established that recombination rate varies across many scales.

Canonical studies in Drosophila demonstrated that recombination rate is heritable, and can itself be selected upon (Chinnici, 1971; Kidwell, 1972). Low to moderate heritability of recombination rate has now been estimated in a large number of additional species including humans, other mammals, insects, and plants (Coop et al., 2008; Dewees, 1975; Dumont et al., 2009; Ebinuma & Yoshitake, 1981; Fledel-Alon et al., 2011; Hadad et al., 1996; Hunter et al., 2016; Johnston et al., 2016, 2018; Jordan et al., 2018; Kawakami et al., 2019; Kidwell, 1972; Sandor et al., 2012; Shaw, 1972). Mapping genetic variants underlying recombination rate variation has identified both local and global modifiers of recombination rate, and the genetic architecture of recombination rate appears to be oligogenic (Dapper & Payseur, 2019; Hunter et al., 2016; Kong et al., 2014; Sandor et al., 2012). Empirical evidence therefore supports the biological plausibility of recombination modifiers distributed across a genome, segregating in populations, and having the potential to increase or decrease recombination rate.

The drift-based model operates under the assumption that the recombination modifier is not under direct selection itself, and that the increase in recombination rate is indirectly favored by the creation of new combinations of alleles. While there is evidence that recombination rate can directly affect fitness, including known effects of sterility of alleles of the recombination modifier Prdm9 in humans and mice (Irie et al., 2009; Mihola et al., 2009; Mukaj et al., 2020), there is also definitive evidence that the generation of new allele combinations by recombination can increase fitness. Comparisons of fitness between populations evolved either asexually or with sexual reproduction and recombination demonstrate that populations that have recombination evolve higher fitness more quickly than populations without recombination (Becks & Agrawal, 2012; Colegrave, 2002; Cooper, 2007; Goddard et al., 2005; Gray & Goddard, 2012; McDonald et al., 2016; Rice & Chippindale, 2001; Zeyl & Bell, 1997). By identifying and testing fitness effects of individual mutations in evolved populations, McDonald et al. showed that increased fitness in sexual populations resulted from recombination unlinking beneficial and deleterious alleles, thereby alleviating clonal interference (McDonald et al., 2016). Using a set of balancer chromosomes that can suppress recombination, selection for bristle number and phototaxis in *Drosophila* demonstrates a significant effect on response to selection as the proportion of the genome allowed to recombine increases (Markow, 1975; McPhee & Robertson, 1970; Rice, 2002). Together, this work demonstrates that the presence, and to some extent the amount, of recombination can be favored and is consistent with theoretical predictions.

Finally, numerous studies have explicitly tested whether recombination rate is increased with directional selection on complex traits that are not known to affect recombination. Selection on traits including flowering time (Harinarayana & Murky, 1971); body weight and litter size (Gorlov et al., 1992); wing length (Palenzona et al., 1972); geotaxis (Korol & Iliadi, 1994); DDT resistance (Flexon & Rodell, 1982); desiccation resistance, hypoxia, and hyperoxia (Aggarwal et al., 2015); and temperature (Gorodetskiĭ et al., 1990; Kohl & Singh, 2018; Winbush & Singh, 2021) have shown significant increases in recombination rate in at least some regions of the genome. While most of these experiments are in *Drosophila* and focus on specific intervals of the genome to score recombination, this body of work does demonstrate that recombination rate can change as a result of directional selection on a wide range of traits over a short time period. The sign and magnitude of change in recombination rate in these studies is consistent with theoretical predictions, which also find variability in the correlated response of recombination to selection, reflecting random effects inherent in the evolution of small populations (Otto & Barton, 2001). However, it remains unclear from these experimental studies if the local effects identified in particular genomic regions extend to increased genome wide recombination rates in selected populations, though some evidence from natural populations suggests this is possible (Samuk et al., 2019).

### Mixed support for the domestication hypothesis of increased recombination

Domestication has been considered an extension of these directional selection laboratory experiments, repeatedly selecting for a number of complex traits, and thus is also hypothesized to increase recombination rate in domesticated species compared to their wild counterparts.

This “domestication hypothesis” was first proposed after Rees and Dale (1974) observed that “bred” lines of grass (*Lolium perenne*) have higher chiasma frequencies than wild lines, followed by Burt and Bell (1987) observing that domesticated species (dog, cat, sheep, goat, horse, cow) have excess chiasma frequencies for their age to maturity compared to other non-domesticated mammals (Burt & Bell, 1987; Jones & Rees, 1966; Otto & Barton, 2001; Rees & Dale, 1974; Ross-Ibarra, 2004).

Direct tests of this domestication hypothesis require estimates of recombination rate in both the domesticated and wild progenitor species. As recombination rate is a challenging phenotype to measure, the task of measuring recombination rate in wild and domesticated organisms is non-trivial (**Box 1**). To date, most studies have been conducted in plants and have yielded mixed results to the domestication hypothesis (**Table 1**). A meta-analysis comparing chiasma frequencies in 26 pairs of wild and domesticated plant pairs supported an increase in recombination rate in domesticated species (Ross-Ibarra, 2004). However, more recent comparisons in rye, barley, and tomato, have shown either no change in inferred recombination rate, or a decrease in recombination rate in domesticated species (Dreissig et al., 2019;

**Table 1.** Comparison of recombination rates between domesticated and wild organisms.

| **Taxon** | **Method** | **Recombination in domestic vs. wild** |  |
| --- | --- | --- | --- |
| Ryegrass | Chiasma | Up | Higher number of crossovers in bred strains compared to wild [1,2] |
| Mammals | Chiasma | Up | Excess number of crossovers in dog, cat, sheep, goat, horse, cow relative to other mammals when corrected for generation length [3] |
| Plants | Chiasma | Up | Higher in domestic compared to wild (analysis included 26 species, including tomato, rice, maize, barley, rye) [4] |
| Cacao | Population | Up | Higher recombination rates in domesticated compared to other wild species [5] |
| Rye | Population | Mixed | No difference in crossover number, but differences in distribution, low-recombining region size, and SC length [6] |
| Tomato | Population | Mixed | Genome wide reduction in effective recombination rate in early domesticated and improved lines; local increases in effective recombination rate in early domesticated [7]. LD highest in domesticated [8] |
| Barley | Population | Mixed | Conservation of recombination landscape with local differences [9] |
| Chicken | ReLERNN | Mixed | Similar or elevated recombination rate in local chicken, similar or decreased recombination rate in industrialized chicken breed [10] |
| Goat | ReLERNN | Mixed | Similar recombination rate in local goat, elevated recombination rate in goat breed [10] |
|  | MLH1 foci | Down | Slight decrease in COs in domesticated compared to wild [11] |
| Soybean | LD decay | Down | Higher LD in domesticated than wild [12] |
| Maize | LD decay | Down | Higher LD, lower rho in domesticated than wild [13, 14] |
| Rice | LD decay | Down | Higher LD in domesticated than wild [15] |
| Dog | MLH1 foci | Down | Slight decrease in COs in domesticated compared to wild [11] |
| Sheep | MLH1 foci | Down | Slight decrease in COs in domesticated compared to wild [11] |
|  | ReLERNN | Down | Similar or decreased recombination rate in local sheep [10] |

[1] Jones and Rees (1966); [2] Rees and Dale (1974); [3] Burt and Bell (1987); [4] Ross-Ibarra (2004); [5] Schwarzkopf et al. (2020); [6] Scheiber et al (2022); [7] Fuentes et al. (2022); [8] Lin et al (2014); [9] Dressing et al (2019); [10] This study; [11] Muñoz-Fuentes et al (2015); [12] Zhou et al. (2015); [13] Hufford et al (2012); [14] Wright et al. (2005); [15] Huang et al. (2010)

Fuentes et al., 2022; Lin et al., 2014; Schreiber et al., 2022). These results contrast the chiasma comparison reported previously for these species (Ross-Ibarra, 2004). Linkage disequilibrium is higher in domesticated soybean, maize, and rice than in their wild counterparts, though this could reflect demography as recombination rate has yet to be examined (Huang et al., 2010; Hufford et al., 2012; Lin et al., 2014; Zhou et al., 2015). Only one study, comparing domesticated cacao to other cacao species, identified a genome wide increase in recombination rate in the domesticated lineage (Schwarzkopf et al., 2020).

A single direct test of the domestication hypothesis has been conducted in animals. A study comparing MLH1 foci (**Box 1**), markers of crossovers, between spermatocytes of dog/wolf, goat/ibex, and sheep/mouflon found a slight increase in crossovers in wild species compared to domesticated animals, refuting the domestication hypothesis (Muñoz-Fuentes et al., 2015).

While this study used small sample sizes (45-184 spermatocytes per species from 2-6 individual male animals), their estimates approximate genetic map length estimates from other studies of domesticated animals, although no such estimates are available for comparison in the wild species. In insects, though no direct comparison was made between domesticated and wild pairs, there is no pattern between high and low recombination rates in domesticated species like the honeybee *Apis mellifera* and silk worm *Bombyx mori* in relation to other non-domesticated species of *Hymenoptera* and *Lepidoptera* (Wilfert et al., 2007).

Motivated by the limited evidence from domesticated animals, we generated genome-wide recombination maps for domesticated and wild populations of chicken, sheep, and goat (**Figure 1**). Recombination rates were estimated independently for each breed/population using ReLERNN and publicly available sequencing data (see Methods) (Adrion et al., 2020; Alberto et al., 2018; Wang et al., 2020). To directly test the domestication hypothesis in chicken, we compared recombination rates between the wild Red Junglefowl (*Gallus gallus spadiceus*), the domesticated but not industrialized local chicken from China (*Gallus gallus domesticus*), and the domesticated Yuanbao chicken breed (*G. g. domesticus*). In support of the domestication hypothesis, we observed that genome-wide recombination rates across local chicken are generally the same or higher than those of the wild species. In contrast, recombination rates across Yuanbao chicken are mixed, with a large portion of recombination rates being the same or lower than in the wild species. Comparison of the wild Iranian Bezoar ibex (*Capra aegagrus*) with the domesticated but not industrialized Moroccan population (*Capra hircus*) and the domesticated Italian Saanen goat breed (*C. hircus*) yielded results more consistently in support of the domestication hypothesis. The majority of recombination rates estimated for the Saanen goat breed were the same or higher than those in the wild species, while the Moroccan population was more mixed, but with most recombination rates exhibiting either no change or an increase. Lastly, we compared a domesticated but not industrialized Moroccan population of sheep (*Ovis aries*) with a wild Asiatic Mouflon (*Ovis orientalis*) population from Iran and found that the domesticated population generally has the same or lower rates of recombination across the genome than the wild species. Though our study is limited in the number of species compared, our data corroborate the collection of studies reviewed here, finding mixed results of the impact of domestication on recombination landscapes.

**Figure 1.**
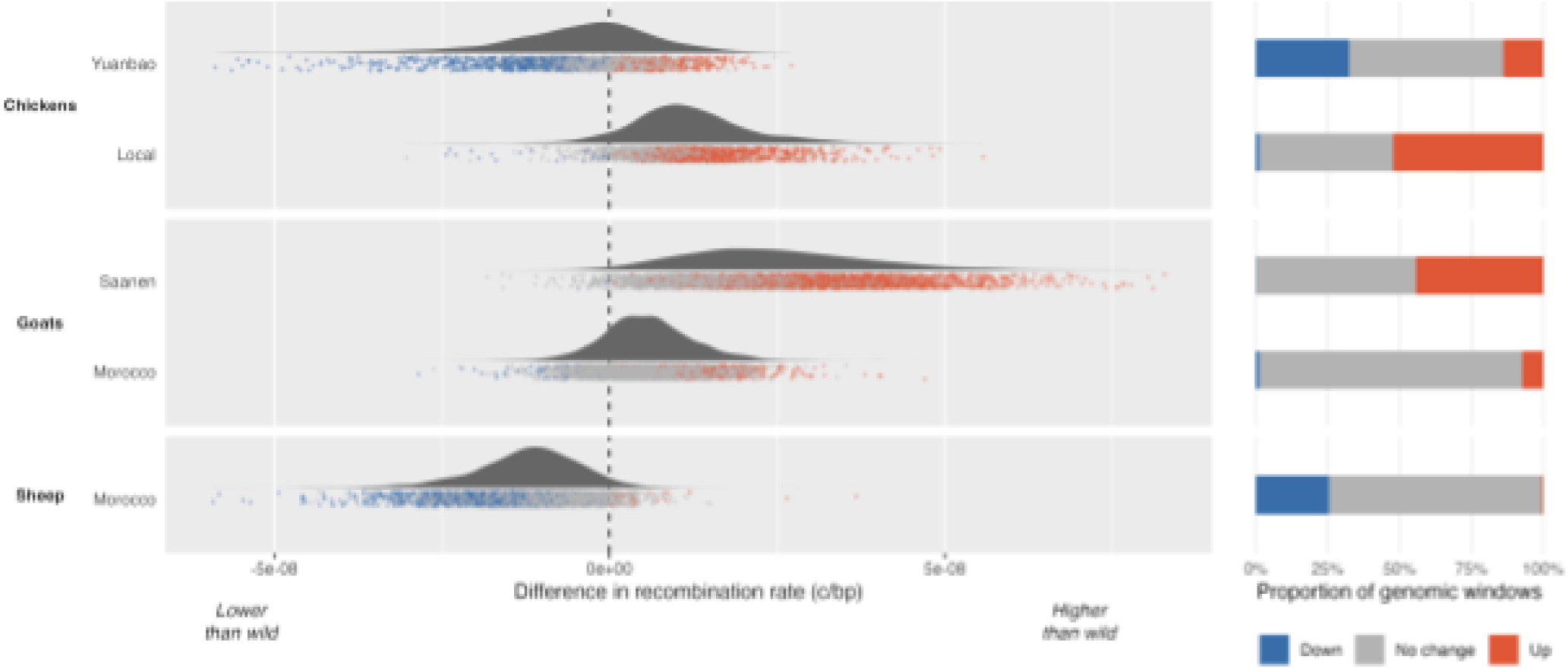
The distributions of recombination rate (in crossovers per base pair) across the genomes of five domesticated lineages do not show a consistent pattern of increase with respect to their corresponding wild relatives. Each point in these distributions represents the difference in recombination rate at a 500-Kb window, with colors indicating the rate was significantly higher (red) or lower (blue) than the wild relative.

While no consistent genome-wide recombination pattern has emerged from plant or animal comparisons, several of these studies have identified local differences in recombination rates between wild and domesticated species. Under the domestication hypothesis, local differences in recombination might be expected in regions containing linked “domestication genes,” genes (or alleles) that were selected on to produce phenotypes in domesticated varieties. For example, local increases in recombination are observed in the early domesticated tomato compared to the wild tomato, including in gene regions that have experienced selective sweeps like resistance (R) genes (Fuentes et al., 2022). As in tomatoes, elevated recombination rates are shifted to distal chromosome regions in domesticated barley compared to wild barley, and are also speculated to result from selection for high recombination in defense response genes during domestication (Dreissig et al., 2019). Comparisons of weedy and domesticated rye show divergence in recombination landscapes; domesticated rye has increased physical regions of low recombination, which may have been selected for by breeders (Schreiber et al., 2022).

Regions containing genes under selection for milk production, coat color, and horn phenotypes in sheep have divergent recombination rates between historical and empirical estimates (Petit et al., 2017). In contrast, no change in recombination was found in domestication genes in wheat or dogs (Danguy des Déserts et al., 2021; Muñoz-Fuentes et al., 2015). While it is clear that there are some differences in recombination between wild and domesticated lineages, one challenge in comparing local shifts in recombination rate between domesticate/wild pairs is that recombination rate is often variable between populations regardless of domestication status (Smukowski & Noor, 2011), thus it is very difficult to untangle an effect stemming from the domestication process itself. However, a pattern of shifts in recombination in regions containing domestication genes is an intriguing avenue for future studies to consider. To match theoretical predictions, a recombination modifier should be linked to loci under selection, but to our knowledge this has not yet been examined.

Domestication genes often show patterns of reduced genetic diversity and differentiation from wild relatives, consistent with selective sweeps. If recombination modifiers are selected to increase recombination rate during domestication, we might expect that some genes involved in recombination show similar patterns of selection to domestication genes. Indeed, several recombination genes show some evidence of selection between teosinte and maize, but do not resemble known maize domestication genes which have very low genetic diversity (Sidhu et al., 2017). Likewise, the *EAS1* ortholog in rye shows evidence of selection in certain domesticated accessions of rye, though not as pronounced as genes involved in traits like seed shattering (Schreiber et al., 2022). *ASY4*, which can affect both number and distribution of crossovers, is associated with divergence in recombination landscapes between landraces of wheat, and missense mutations in the *FIGL-1* ortholog in cacao are associated with increases in recombination in domesticated cacao (Danguy des Déserts et al., 2021; Schwarzkopf et al., 2020). With an increasing number of datasets on between populations/species recombination rate landscapes and a list of candidate recombination genes, future studies may increasingly be able to connect suggestive recombination rate modifiers to the evolution of recombination rate.

### Rectifying theory and data

Taken together, no clear pattern emerges linking domestication to an increase in recombination rates in plants or animals. How do we make sense of differences in predictions from theory, and between experimental studies that point to conflicting conclusions? First, we must consider how methodologies used to estimate recombination rate might contribute to conclusions (**Box 1**, see review of methods (Peñalba & Wolf, 2020)). Early work estimating recombination generally used chiasma counts, which can be challenging as a proxy for crossover counts. The condensed chromosomes at entry to metaphase, the stage at which true chiasmata can be identified, causes detection to be difficult, and a variety of other variables can lead to an overestimation of chiasmata counts (Hultén, 1974; Muñoz-Fuentes et al., 2015; Wada & Imai, 1995). The field now typically relies on MLH1 foci and other molecular markers to identify crossovers, and this could account for discrepancies between conclusions from early studies and more recent linkage disequilibrium (LD) based recombination maps. The vast majority of data has used LD-based methods to infer the historical averaged population recombination rate (⍴) for domesticated and wild pairs, but using LD to estimate recombination rate can come with its own challenges. A reduction in nucleotide diversity caused by selective sweeps, bottlenecks, and/or inbreeding, all common processes during domestication, can obscure estimating the true recombination rate (r) from ⍴. Simulations have demonstrated that a reduction in nucleotide diversity caused by a recent bottleneck can lead to an underestimation of the true rate of recombination (Chan et al., 2012), and the effects of selection, demography, and gene flow on detecting recombination rate may be more acute at fine scales (Dapper & Payseur, 2018; Samuk & Noor, 2022). That being said, strong correlations between cross-based and LD-based maps in a wide range of organisms, including many of the domesticated lineages covered in this review, suggests that this is not a major source of bias, at least at broad genomic scales. The method we utilized to estimate recombination in chicken, goat, and sheep is robust to various demographic models (Adrion et al., 2020), but demography could impact our results as well. Disentangling effects of a change in effective population size and recombination rate in populations and in regions of the genome that have experienced strong selective pressure may deserve further attention.

Second, the point in time at which domesticated and wild species genomes or meioses are sampled may influence our interpretation of recombination rate. It is possible that the recombination landscapes of both wild and domesticated species have changed in the centuries following their split, and thus inferring recombination in extant organisms fails to capture true changes in recombination between wild progenitors and domesticated descendants. Similarly, the phase in the domestication process (e.g., the spectrum from wild organisms to landraces/village animals to breeds/improved lines) may be important. A decrease in recombination rate is favored when high fitness allele combinations are present on the same haplotype (Barton, 1995; Barton & Charlesworth, 1998; Otto & Lenormand, 2002). An increase in recombination may occur during domestication, but it may be ephemeral, as breeders select against recombinants that break up desired trait combinations, and/or an increase in homozygosity reduces the effect of recombination. This may help explain patterns in tomato, where local increases in recombination are detected in early domesticated tomato from wild tomato, but are highly reduced in the improved tomato line (Fuentes et al., 2022), and in chicken, where the local chicken has increases in genome wide recombination rate compared to junglefowl, but the Yuanbao breed has a similar or reduced genome wide recombination rate. However, no such pattern is detected in goats, where both the local domesticated goat and the Saanen breed have increased recombination rates compared to the wild species.

Finally, local changes in recombination rate between lines/breeds have been identified in a wide array of domesticated species (Bauer et al., 2013; Brazier & Glémin, 2022; Danguy des Déserts et al., 2021; Groenen et al., 2009; Ma et al., 2015; Petit et al., 2017; Q et al., 2016; Weng et al., 2019). This variation between accessions/breeds suggests a capacity for rapid changes in recombination rate, but also makes it challenging to ascribe these changes to domestication. Furthermore, which population is sampled from the wild and domesticated lineages for comparison could lead to different interpretations. For example, it is feasible that one breed could have increased regions of recombination rate compared to a wild progenitor, but a related breed could have similar or decreased recombination rate. Our results of recombination rate landscapes in goat differ from those reported using MLH1 markers, which found more crossovers in the wild Spanish ibex compared to local goats (Muñoz-Fuentes et al., 2015), but our study used different populations/breeds of both domesticated and wild goats. Our results for sheep follow the same pattern as the estimates from MLH1 foci, but again, we used different populations of both domesticated and wild species. Elevated local differences in recombination in domesticated organisms could indeed support the domestication hypothesis, but variation between populations of both wild and domesticated organisms must be taken into account.

## Conclusions

Experimental studies in the lab often show an effect of directional selection on altering recombination rates. The more inconsistent patterns in recombination rate in domesticated organisms perhaps highlight that the domestication process is complex, reflecting other forces at work. Future studies could return to more controlled studies in the lab or field, harnessing experimental evolution with directional selection and leveraging sequenced based methods to measure genome wide recombination and connect this with the genetic architecture of loci under selection. This type of experimental work coupled with simulations, including further exploration of recombination rate as a complex trait (Drury et al., 2023), could further elucidate the strength of selection, number of generations of selection required, genetic architecture of traits under selection, local vs. global effects of modifiers, number of modifiers, and effect size to the evolution of recombination rate.

Breeders have begun pursuing an exciting avenue leveraging modifiers to tune recombination rates in domesticated species for crop improvement (Epstein et al., 2023; Fayos et al., 2019; Kuo et al., 2021; Taagen et al., 2020). Variants or mutants of recombination modifiers *HEI10*, *FANCM*, *FIGL1*, *FLIP*, and *RECQ4* can increase crossover frequency up to 8x in Arabidopsis (Fernandes et al., 2018; Serra et al., 2018; Ziolkowski et al., 2017). Preliminary attempts to test some of these genes in crop species have been successful, with *recq4* mutants increasing crossover frequency 3-fold in rice, pea, and tomato (de Maagd et al., 2020; Mieulet et al., 2018). Altering epigenetic patterns like methylation has been shown to change the number and distribution of crossovers in Arabidopsis and maize, and more specific targeting of crossovers to particular targets is an active area of exploration. While additional work is needed to assess impacts on fitness and other traits before this approach is likely to be utilized as a strategy, simulations have shown that modifying the crossover number of crops could improve genetic gain and maintain more genetic variance (Epstein et al., 2023). Apart from the applied use of these recombination modifiers, the ability to influence the frequency and distribution of crossovers is an exciting opportunity for further exploration of the evolution of recombination rate in experimental work.

## Methods

### Data Preparation

To better understand how domestication shapes recombination, we generated new comparisons of recombination rate for sheep/mouflon, goat/bezoar, and chicken/junglefowl. We collected data from publicly available VCFs (Alberto et al., 2018; Wang et al., 2020). We selected individuals from populations in the same geographical area with shared population structure and used bcftools v1.13.0 to extract individual VCFs for each breed or population (n=10 samples when available, **Table S1**). For Red Junglefowl, Yuanbao chicken, and Local chicken, we pulled 10 samples each. For Bezoar, we pulled all samples originating from Azerbaijan (n=8), for Moroccan domesticated goats we pulled 10 samples at random (n=10), and for Italian Saanen goats we used all available samples (n=5). For Mouflon and Moroccan domesticated sheep, we pulled 10 samples at random from each population (n=10). For each population of interest, we used bcftools to filter for autosomes and biallelic SNPs. We then used vcffixup v1.0.3 to recalculate allele frequencies and filter for sites with MAF greater than 0.05 and less than 0.95 (Garrison et al., 2022). Additional filtering was applied for chicken populations to remove chromosomes with less than 250 SNPs. Per ReLERNN requirements, we generated a BED-formatted file of chromosome positions using the UCSC Genome Browser fetchChromSizes script and the reference genomes used to generate the original VCFs (Nassar et al., 2023).

To mask structural variants between wild and domesticated genomes, each pair of wild and domesticated chromosome-level genome assemblies were aligned to each other using minimap2 with default parameters (Li, 2018). The genome assemblies Gallus_gallus-4.0 (chicken), Oar_v3.1 (sheep), and CHIR_1.0 (goat) were used as the references and GRCg6a (red jungle fowl), CAU_Oori_1.0 (mouflon; GCA_014523465.1,), and CapAeg_1.0 (bezoar) were used as queries (Bellott et al., 2017; Dong et al., 2013, 2015; Hillier et al., 2004; Jiang et al., 2014). The SAM file generated by minimap2 was then used as input for SyRI to annotate regions that contain structural variants (Goel et al., 2019). SyRI outputs a TSV that was then formatted as a BED file for ReLERNN, retaining non-overlapping genomic coordinates at which structural variants were inferred using the bedtools ‘merge’ function (Quinlan & Hall, 2010).

### Estimating recombination rates with ReLERNN

As input for ReLERNN (Adrion et al., 2020), we provided the processed VCFs for each population, the BED file of chromosome positions, and an accessibility mask file. The upperRhoThetaRatio or URTR parameter sets an upper bound for recombination rate estimates based on the ratio of rho and theta. For chicken, we used URTR=40, based on a mutation rate of 2.0e-9 bp/generation and recombination rate of 8 cM/Mb (Bird et al., 2020; Groenen et al., 2009; Shi et al., 2023). For sheep and goat, we used URTR=4 based on a mutation rate of 2.5e-8 bp/generation and a generation length of 2 years and recombination rates of (Alberto et al., 2018; Johnston et al., 2016; Petit et al., 2017). For ReLERNN_TRAIN, we used the default nEpochs = 1000, which ran until the calculated training loss reached the early stopping threshold set by ReLERNN. For chicken, ReLERNN met the early stopping threshold after 394, 375, and 324 epochs for Red Junglefowl, Local chicken, and Yuanbao chicken, respectively. For goats, ReLERNN met the early stopping threshold after 315, 343, and 302 epochs for Bezoar goats, Moroccan goats, and Saanen goats, respectively. For sheep, ReLERNN met the early stopping threshold after 280 and 277 epochs for Mouflon sheep and Moroccan sheep, respectively. Addition of the bed-formatted accessibility mask resulted in 2.8%, 2.7%, and 2.8% of the genome being masked for structural variants in Red Junglefowl, Local chicken, and Yuanbao chicken, respectively. 21%, 21.1%, and 21% of the genome was masked for structural variants in Bezoar goats, Moroccan goats, and Saanen goats, respectively. 9.9% of the genome was masked for structural variants in both Mouflon sheep and Moroccan sheep. All other parameters used default values recommended by ReLERNN (https://github.com/kr-colab/ReLERNN).

### Comparing recombination rate estimates

For each species pair, we carried out comparisons of recombination rates in non-overlapping 500-kb windows (using R version 4.4; (Wickham et al., 2019)). These windows result from averaging the rates inferred by ReLERNN in the corresponding regions (which varied in size, typically around 120 kb), weighted by the number of bases that overlap the focal 500-kb region. To assess the statistical significance of the differences in recombination, we used the inferred 95% confidence intervals at the ReLERNN window corresponding to the start of the 500-kb region. We classified windows as having significant changes in recombination (coded as ‘up’ or ‘down’ in Figure 1) when the intervals of the two compared lineages did not overlap.

## Data and Code availability

The scripts for these analyses, a more detailed description of our data preparation steps, and the final output from ReLERNN are available in the Supplement.

## Box 1

### Methods of estimating recombination rate

#### Cytology based methods

imaging based methods that count crossovers in individual cells going through meiosis. Crossovers are approximated by counting chiasmata, the physical link between chromatids of homologous chromosomes where genetic material is exchanged, or counting foci based on immunolocalization of proteins that mark mature crossovers (MLH1 is commonly used in mammals). Provides coarse resolution of genome-wide recombination rate.

#### Cross/pedigree-based genotyping

Utilizes a cross between two genetically distinct parents or pedigrees and molecular markers (microsatellites, SNPs, etc) to estimate present day recombination rate (cM/Mb). Can be sex-specific. Resolution depends on marker density and number of individuals, but can provide fine scale estimates.

#### Linkage disequilibrium (LD) based methods

Computational methods that use population sequencing data to infer the sex-averaged historical population recombination rate, ⍴ (4N_e_r).

#### Machine learning based methods

Computational method that uses population sequencing or pooled sequencing data and recurrent neural networks (Adrion et al., 2020). This algorithm likely learns from the ratio of recombination and mutation rates in simulations to make its inferences.

## Acknowledgements

We thank Daniel Lucio for assistance in installing and running ReLERNN on the NCSU Bioinformatics Research Center computing cluster. This work was supported by NIH R35 GM142849 to C.S.H. and NIH R35 GM147107 to R.F.G.

## Notes

### Competing Interest Statement

The authors have declared no competing interest.

